# Fluorogenic, sub-single-turnover monitoring of enzymatic reactions involving NAD(P)H provides a generalised platform for directed ultrahigh throughput evolution of biocatalysts in microdroplets

**DOI:** 10.1101/2023.11.22.568356

**Authors:** Matthew Penner, Oskar James Klein, Maximilian Gantz, Friederike E. H. Nintzel, Anne-Cathrin Prowald, Sally Boss, Paul Barker, Paul Dupree, Florian Hollfelder

## Abstract

Enzyme engineering and discovery are crucial for a future sustainable bioeconomy. Harvesting new biocatalysts from large libraries through directed evolution or functional metagenomics requires accessible, rapid assays. Ultra-high throughput screening formats often require optical readouts, leading to the use of model substrates that may misreport target activity and necessitate bespoke synthesis. This is a particular challenge when screening glycosyl hydrolases, which leverage molecular recognition beyond the target glycosidic bond, so that complex chemical synthesis would have to be deployed to build a fluoro- or chromogenic substrate. In contrast, coupled assays represent a modular ‘plug-and-play’ system: any enzyme-substrate pairing can be investigated, provided the reaction can produce a common intermediate which links the catalytic reaction to a detection cascade readout. Here, we establish a detection cascade producing a fluorescent readout in response to NAD(P)H via glutathione reductase and a subsequent thiol-mediated uncaging reaction, with a low nanomolar detection limit in plates. Further scaling down to microfluidic droplet screening is possible: the fluorophore is leakage-free and we report a three orders of magnitude improved sensitivity compared to absorbance-based systems, so that less than one turnover per enzyme molecule expressed from a single cell is detectable. Our approach enables the use of non-fluorogenic substrates in droplet-based enrichments, with applicability in screening for glycosyl hydrolases and imine reductases (IREDs). To demonstrate the assay’s readiness for combinatorial experiments, one round of directed evolution was performed to select a glycosidase processing a natural substrate, beechwood xylan, with improved kinetic parameters from a pool of >10^6^ mutagenized sequences.

## Introduction

Enzyme engineering campaigns rely on functional screening for the discovery of starting points and subsequent directed evolution^1^. In light of the increasing demand for biocatalysts in the transition towards a sustainable bioeconomy^2^, sensitive assays compatible with ultrahigh-throughput screening formats are crucial for overcoming the challenges arising from finding rare functions in the proverbial vastness of sequence space^3,4^ in a time-efficient manner^5,6^. However, spectroscopic assays routinely used for enzyme screening rely on bespoke model substrates containing aromatic fluorophores or chromophores released as leaving groups upon catalysis^4,6–12^, but these model substrates rarely emulate the recognition features of many target reactions. Moreover, following directed evolution’s basic law “*you get what you screen for*” adaptation of a target enzyme to a model substrate can result in minimal improvements to the desired reaction of a natural substrate^13,14^. Conversely, by coupling the enzymatic reaction to a reporter reaction by means of a common co-substrate, such as NAD(P)H, substrates that are neither fluorogenic nor chromogenic themselves^15,16^ may be assayed using fluorescent or absorbance based techniques. As any enzymatic reaction dependent of the common co-substrate can be coupled to such an assay, these are versatile (“plug-and-play”) with little change in setup required between different classes of enzymes. Beyond their use in enzyme discovery and engineering, these assays may also function as analytical tools for the quantitative detection of small molecules within complex mixtures^17,18^ by virtue of the intrinsic specificity afforded by enzymes. In such a case, the engineerability of reporter enzymes enables the creation of tailored sensors for a desired analyte. In this study we introduce the novel fluorogenic dinitrophenyl sulfonyl coumarin probe (**S**_N_**A**r probe **F**or **R**apid **A**ssay of **N**ADPH, *SAFRAN*) in a coupled assay format for quantification of NADH and NADPH-dependent enzymatic activity at ultra-high throughput. We demonstrate SAFRAN’s ability to assay both NADH-forming and NADPH-consuming reactions and exemplify coupled droplet-based selections for two representative enzyme classes: (i) a glycosyl hydrolase linked to a sugar dehydrogenase as a sensor enzyme and an (ii) an imine reductase (IRED) that consumes NADPH for the synthesis of chiral amines from ketones.

The reduction of NAD(P)^+^ to NAD(P)H itself can be monitored by an increase in absorbance at 340 nm, or *vice versa* for reductive enzymes. This colorimetric change lies at the heart of many small-molecule quantification kits, such as those sold commercially for monosaccharide detection^19^. However, the low molar extinction coefficient of NADH imposes a de facto detection limit of 10 µM in plates using direct absorbance of the cofactor^20^. Previous strategies to address these limits have focused on coupling NADH reduction to tetrazolium dye reduction^20–22^, boosting the measured molar absorption coefficient and improving the signal of the assay 3-fold. Despite these advances, miniaturized ultrahigh-throughput experiments in microfluidic droplets still suffer from a substantial absorbance detection limit of around 10 µM even while using costly dyes^5,6,20^. Improvements in sensitivity can be achieved using fluorescence assays, as they do not rely on light fully traversing the analyte solution and detect at a wavelength different to the source^23^. While NAD(P)H is itself a fluorophore, its quantum yield is low (2% in aqueous solution, leads to no major sensitivity improvement from using fluorescence compared to absorbance in plate format)^24,25^, and its fluorescence lifetime as well as excitation/emission maxima are dependent on protein binding^26,27^, complicating assays in complex mixtures such as cell lysate^28,29^. Coupling a fluorogenic reaction to NAD(P)H is therefore a compelling alternative, boosting the sensitivity of fluorescent detection while overcoming the limitations of molecular recognition. However, past efforts to do this have been hindered by droplet-droplet leakage of the fluorescent product^30^. We achieve this by an enzymatic cascade that detects NAD(P)H and results in the activation of SAFRAN, a pro-fluorescent probe designed to be leakage-free in microfluidic droplets.

The improvement of glycosyl hydrolases is of direct interest for sustainable biocatalysis and members of the family GH115 are particularly promising in this context as they debranch the hemicellulose xylan, removing internal glucuronic acid residues^31^. These functional groups are crucial for the recalcitrance of biomass to enzymatic degradation^32^. Enzymes to remove these linkages ever more efficiently promise clean ways to utilize the plant cell wall^33^. However, low activity and poor expression have held back industrial applications of enzyme candidates^34^, in which cocktail formulations must be finely tuned, while increased enzyme loadings alone are frequently insufficient for optimal sacchrification^35,36^. Therefore, it has been hypothesised that engineering of members of the GH115 family to be more active towards a broader range of substrates will unlock the promise of this family for biocatalytic applications^35^. High-throughput assay of the glycosidase reaction has, however, been hampered by the molecular recognition of the substrate routinely extending far beyond the target glycosidic bond itself^37,38^. Thus, when model substrates lacking specificity-defining features were used to provide an optical readout in directed evolution campaigns after cleavage, the kinetic gains shown by the target enzyme did not translate well to the actual target substrate, as shown in Table S1, creating demand for methods that permit glycosidase screening against natural substrates^39–41^. We achieve such assay by linking the release of glucuronic acid residues with NADH formation, and further to SAFRAN, *via* a specific dehydrogenase acting as a reporter (**Figure 1**). As an abundance of selective, NAD(P)H-dependent oxidoreductases have been described^42^, this methodology can be easily expanded to detect a wide range of functional groups and small molecules.

**Figure 1:**
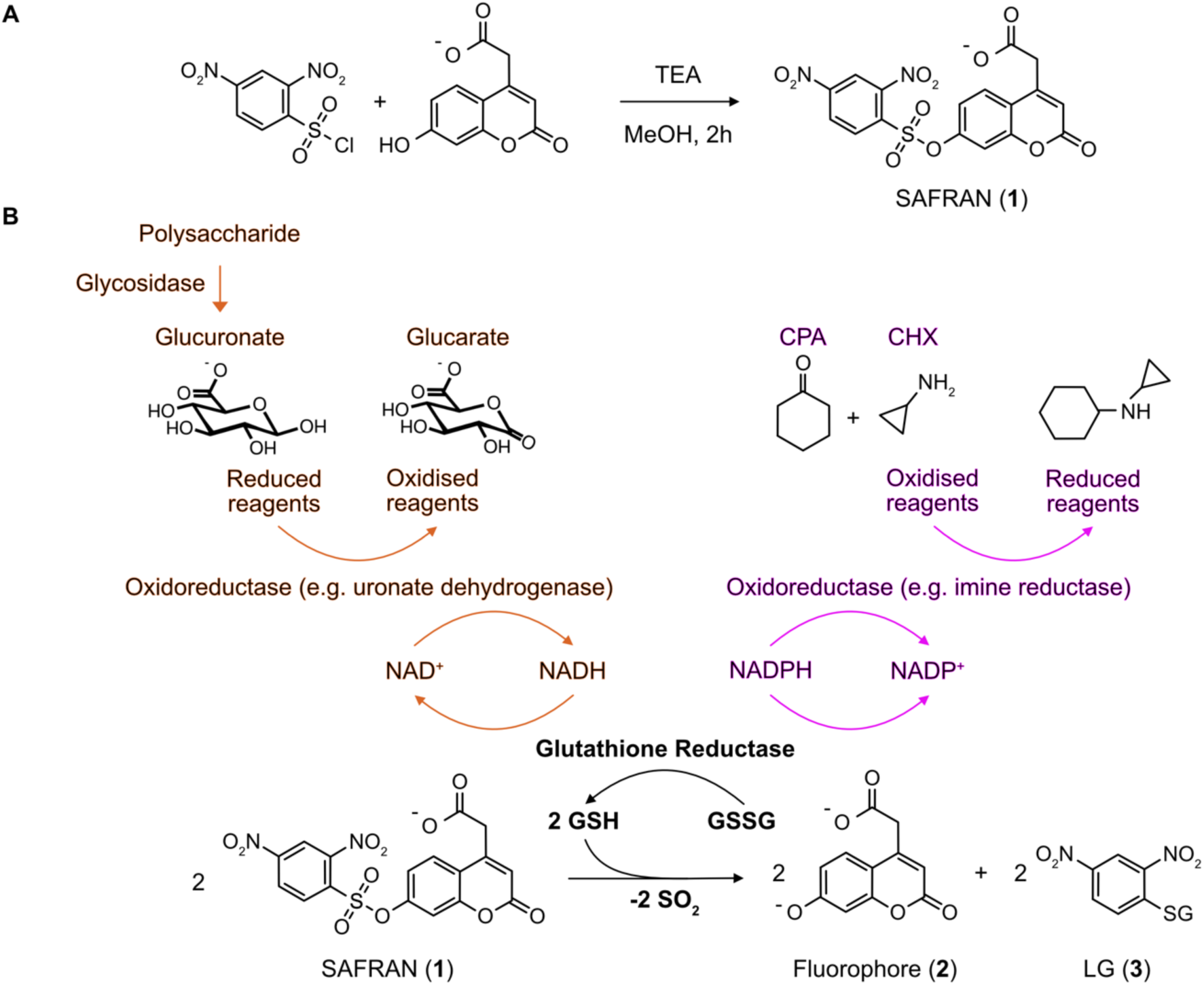
SAFRAN synthesis and coupled assay scheme. (A) Synthesis of SAFRAN **1** from commercially available components; (B) An analyte released by an *upstream* glycosidase is oxidized (orange) by a specific *downstream* oxidoreductase, producing or consuming the NAD(P)H that serves as substrate for glutathione reductase. Reduced glutathione (GSH) spontaneously reacts with SAFRAN **1** forming the coumarin fluorophore product **2** (and the GSH adduct **3**). Alternatively, a primary reaction can be detected that modulates NADP^+^/NADPH interconversion (purple). In this scheme, an imine reductase consumes NADPH, which can then be titrated by the *downstream* detection cascade.

For an application of this new assay in the reverse direction (i.e. for NADH consumption rather than formation), we chose to investigate an enzymatically catalysed imine reduction reaction, in which NAD(P)H is depleted in a C-N bond forming reaction, for the specific enzyme in question, using the phosphorylated cofactor NADPH instead of NADH. Imine reductases (IREDs) are widely used to access chiral amines, a crucial functionality found abundantly in pharmacologically interesting compounds^43^. As most IREDs require engineering before becoming industrially viable and most reactions do not involve strongly fluorogenic compounds, there is a demand for rapid, sensitive assays in the field. As this reaction directly consumes NAD(P)H, SAFRAN can be applied to back-titrate and quantify NAD(P)H depletion in picodroplets, replacing the less-sensitive tetrazolium dye method^44^.

We benchmark the application of SAFRAN by quantification of NAD(P)H concentrations and, specifically for glycosyl hydrolases, of monosaccharides at ultrahigh-throughput while still matching the detection limit of High-Performance Anion-Exchange Chromatography with Pulsed Amperometric Detection (HPAEC-PAD), the acknowledged gold-standard column-based method for monosaccharide detection^32,45,46^. Compared to previously reported coupled assays for NAD(P)H detection in droplets with a low micromolar limit of detection^20^, our system provides a 300-fold improvement in sensitivity and maintains the integrity of the product readout over multiple days, enabling extended incubation times.

Having quantified the scope of this assay, we demonstrate its utility in ultrahigh throughput screening in microfluidic droplets by substantial enrichments in mock library selections (from defined plasmid mixtures) for both example reactions, glycosidases and IREDs, in forward and reverse assay directions. Finally, we apply the SAFRAN cascade in actual directed evolution, starting from a library of mutant GH115 glycosidases and screen >10^6^ members for glucuronoxylan-debranching activity. We identify and recover an abundance of mutants with improved lysate activity and, after purification of five of the mutants, identified a sextuple mutant with a 2-fold increase in *k_c_*_at_. Based on these findings, we propose that the SAFRAN coupled assay represents a widely applicable method for enzyme discovery, directed evolution and small molecule quantification in picodroplets.

## Results and Discussion

### A NAD(P)H-based detection cascade based on thiol-dependent uncaging of Safran

To couple NAD(P)H production with turn-on fluorescence, we designed a cascade linking the emergence of NAD(P)H to the formation of free thiols *via* enzymatic glutathione reduction. The free thiols subsequently uncage a synthesised fluorogenic probe (SAFRAN) in a rapid and irreversible S_N_Ar reaction with a dinitrophenol group that was quenching coumarin fluorescence^47–50^. Synthesis of SAFRAN **1** was readily achieved by condensation of 7- hydroxycoumarinyl-4-acetic acid **2** with 2,4-dinitrobenzenesulfonyl chloride in one step from commercially available compounds at 76% yield without the need for purification by chromatography (**Figure 1A**). High sensitivity is ensured with a favourable reaction stoichiometry – two equivalents of reduced glutathione are produced for each equivalent of NAD(P)H consumed. Any NAD(P)H-producing enzyme could in principle be added to this cascade to elicit a fluorogenic response (**Figure 1B**). The cascade reagents are cheap and generally accessible – NAD(P)H is re-oxidised by the commercially-available glutathione reductase, so the expensive NAD(P)^+^ cofactor only needs to be present in catalytic quantities.

The design of SAFRAN was focused around a strongly fluorescent 7-hydroxycoumarin core – as coumarin derivatization is robust and well established, the probe could be readily functionalised^51,52^. Leakage is a well-documented challenge in microfluidic assay development, preventing the use of the typically large hydrophobic and aromatic probes for all but the fastest reactions^53–57^. This poses a hitherto unsolved challenge for fluorogenic NAD(P)H quantification as resorufin, the sole published fluorogenic probe for NAD(P)H detection^54,58^, is hydrophobic and consequently only compatible with very short reaction times in microfluidic droplets^59^. To minimize leakage of SAFRAN and its fluorophore product **2**, a carboxylic acid group was incorporated to impart a negative charge on the coumarin scaffold at physiological pH, enabling application in droplet microfluidics with minimal leakage.

As the fluorescence of the umbelliferone structure is dependent on its protonation state (determined by the pH of the reaction medium^60^), we obtained fluorescence excitation and emission spectra of the 7-hydroxycoumarin fluorophore under different pH conditions (**Figure S1**). Under physiological to slightly basic conditions, the highest signal intensity was achieved with an excitation wavelength of 380 nm coupled to an emission wavelength of 460 nm.

### Detecting picomoles of monosaccharides in high throughput

To determine the sensitivity of SAFRAN **1** to reduced glutathione (GSH), GSH was added across a concentration range (25 nM to 25 µM), eliciting a linear response over three orders of magnitude after ninety minutes of incubation (**Figure 2A, S2**). Formation of the phenylated glutathione product **3** and release of the coumarin fluorophore **2** were confirmed *via* mass spectroscopy (**Figure S3**). Subsequently, we demonstrated that glutathione reductase couples the presence of NADH to the S_N_Ar reaction and thus fluorogenic signal (**Figure 2B**), with a detection limit of 30 nM NADH (corresponding to an amount of 3 pmol NADH, p = 0.001, **Figure 2B, S4**) in plates. Taking advantage of the bi-specificity of glutathione reductase for both nicotinamide cofactors NADH and NADPH^61^, we furthermore show the applicability of the SAFRAN cascade for quantification of NADPH in plate format (**Figure S5**).

**Figure 2:**
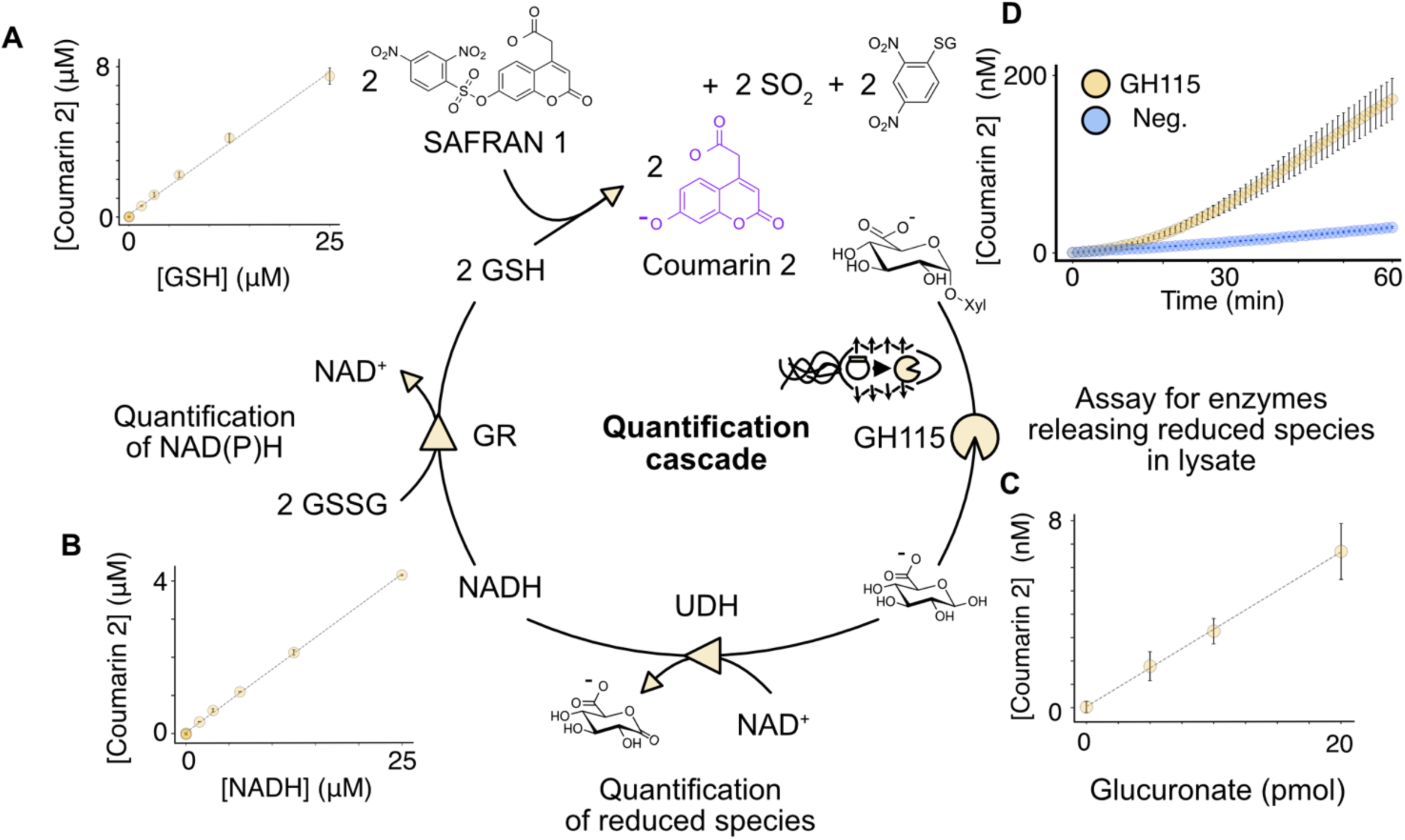
A versatile coupled assay enables detection of trace amounts of monosaccharides in plate format. (A) Unless otherwise stated, all reactions were measured in triplicate at 23 °C, in 200 mM Tris-HCl (pH 7.0), excited at 380 nm and its emission measured at 460 nm. Quantification of fluorescence emission after reaction completion (after 90 minutes) shows a linear response to reduced glutathione addition. The limit of detection was 50 nM, as defined by a signal significantly above baseline (p = 0.04, Welsh’s one-sided t-test, n = 3 per sample) (B) The S_N_Ar reaction was coupled to glutathione reductase to consume NADH and product quantification was achieved based on a linear calibration curve, in this case using fluorescence emission after 90 minutes of incubation. The limit of detection was 30 nM (p = 0.001, Welsh’s one-sided t-test, n = 3 per sample) (C) The cascade can be further coupled to a monosaccharide dehydrogenase (uronate dehydrogenase) facilitating accurate quantification of glucuronate concentration with a detection limit of 5 pmol per well, or 25 nM (p = 0.02, Welsh’s one-sided t-test, n = 3 per sample). Superior sensitivity was achieved by using 200 mM Tris-HCl (pH 8.0), and over one hour (the measurement time point) the reaction had reached completion. (D) Time-course of fluorescence emission of GH115-expressing cells with glucuronoxylan substrate and assay components, compared to cells expressing the control enzyme *Sr*IRED, demonstrating that assay can be applied in lysate for quantification of the degradation of complex saccharides or polymers. Xyl = xylan.

To demonstrate the use of the SAFRAN assay as an analytical tool with high sensitivity, we set out to show its potential for improving monosaccharide detection limits in high throughput. Monosaccharide liberation is often associated with the breakdown of complex sugars. However, the most sensitive monosaccharide quantification currently, HPAEC-PAD, requires minutes per single assay, very clean samples^62^ and large amounts of costly and sometimes unstable standards^63^. In contrast, high throughput plate-based detection of monosaccharides has sensitivity in the micromolar range at best and can suffer from low selectivity for the monosaccharide of interest^64^. Compared to HPLC-based systems or direct chemical detection, dehydrogenase enzymes are engineerable^65^ and a suite of these enzymes are already available as coupling agents^42^, so that tailored sensors for any analyte or application can be created.

Using the coupled assay in plate format and selecting uronate dehydrogenase as a representative coupling enzyme, we were able to detect glucuronic acid as a linear signal with a detection limit of 25 nM (p = 0.02, **Figures 2C and S6**), representing only 5 picomoles of analyte per well. This is a >100-fold improvement on the best commercially available plate-based glucuronic acid quantification method^19^ and comparable to the 12.5 picomole/sample limit of quantification of HPAEC-PAD^66^. In addition to fluorescence end-point measurements, monosaccharides could be also quantified within minutes of reaction initiation using a maximum rate method, enabling shorter analysis times (**Figure S6**). This indicates that the method is a rapid, accessible and derivatisation-free alternative to column-based methods for trace monosaccharide quantification, promising to reduce the equipment and reagent costs involved in sensitive monosaccharide detection.

Small molecule quantification is a widely-used analytical method for quantifying polymer degradation^58^. Upon confirming that the SAFRAN assay was well-suited for quantifying small molecules, we set out to test the capacity of the cascade to quantify the activity of glycosidases that debranch hemicellulose polysaccharides, using *E. coli* as the enzyme expression host. In a cellular context, detection cascades risk suffering interference, both optically and chemically. To test the theoretical limits of detection of the cascade in *E. coli* lysate that may be in place due to optical interference, we diluted coumarin 2 in a working concentration of *E. coli* lysate and determined the limits of detection to be 8 nM of coumarin 2 (p = 0.0001, **Figure S7**).

We then set out to evaluate whether chemical interference in the cascade would preclude screening for catalysis in an *E. coli* host. If background activities in the screening host are too high, excessive background signal can make quantification of target activity impossible. This would reduce the capacity of the cascade to detect enzyme activity in cell lysate, thereby limiting its use for high-throughput enzyme screening. To test the signal to noise ratio of this cascade, we incubated *E. coli* pre-induced for expression of a glucuronoxylan-debranching enzyme (*AxyAgu*115A) with lysis agent, cascade components and glucuronoxylan substrate over an hour-long window. This revealed that, following a brief lag-phase, fluorescence intensity increased up to 4-fold above the negative lysate control within an hour, following glucuronic acid release from the polymer (**Figure 2D**). The ability to use our coupled assay for screening of enzymes in *E. coli* lysate integrates it into a preferred format for plate and droplet-based enzyme engineering and metagenomic screening campaigns^67^.

To demonstrate the broad applicability of the SAFRAN cascade we also explored its ability to report on a NADPH-dependent reduction reaction, thus highlighting both the tolerance of glutathione reductase for phosphorylated and non-phosphorylated nicotinamide and the capability to use this assay for back-titrations. To this end, the condensation reaction between cyclopropylamine (CPA) and cyclohexanone (CHX) *via* reductive amination, catalysed by an imine reductase from *Streptosporangium roseum* (*Sr*IRED) (**Figure 1B**), was examined. After reductive amination using lysate containing *Sr*IRED, the cascade was added to back-titrate IRED activity showing a clear time-dependent signal (**Figure S5**).

### Detection of oxidoreductases in picodroplets using SAFRAN

Much larger numbers of protein variants can be screened in an attractive ultrahigh throughput format where water-in-oil emulsion droplets act as reaction compartments made and handled in microfluidic devices^6^. However, this format has specific limitations that must be overcome, e.g. the leakage of aromatic, hydrophobic fluorescent probes between droplets, likely due to enhanced oil-probe hydrophobic interactions^56^. This leakage results in a loss of signal over time, shortening maximum droplet incubation times and thus precluding the detection of enzymes that are poorly expressed or have low activity. Therefore, SAFRAN was designed to carry a net negative charge at physiological pH in the form of a carboxylic acid moiety. Indeed, fluorescence microscopy imaging of a mixed population of droplets, containing either buffer only or 100 µM coumarin **2**, showed no equilibration between droplets after 24 hours (**Figure S8**). Further, incubation of a mixed population of droplets containing 25 μM and 50 μM of fluorophore revealed no detectable leakage for up to five days (**Figure 3A**) when measured using Fluorescence-Activated Droplet Sorting (FADS). Measuring the same droplets again at day 16 showed that the fluorescence of the two populations had begun to converge (by 73% of their original fluorescence). However, the individual populations could still be distinguished with only 3% overlap. This indicates a very slow rate of leakage, perhaps inevitable due to surfactant-mediated transfer between droplets^68,69^. These measurements indicate that incubations of up to at least five days should be feasible without facing challenges with leakage. This stands in contrast to the NAD(P)H probe resorufin – fluorescence microscopy in a previous study revealed near complete equilibration of droplets with and without fluorophore in only 6 hours^59^, suggesting SAFRAN achieves at least 64-fold better signal retention over this critical window used in picodroplet screening for medium-to-slow reactions (Table S2)^7,9,70^.

**Figure 3:**
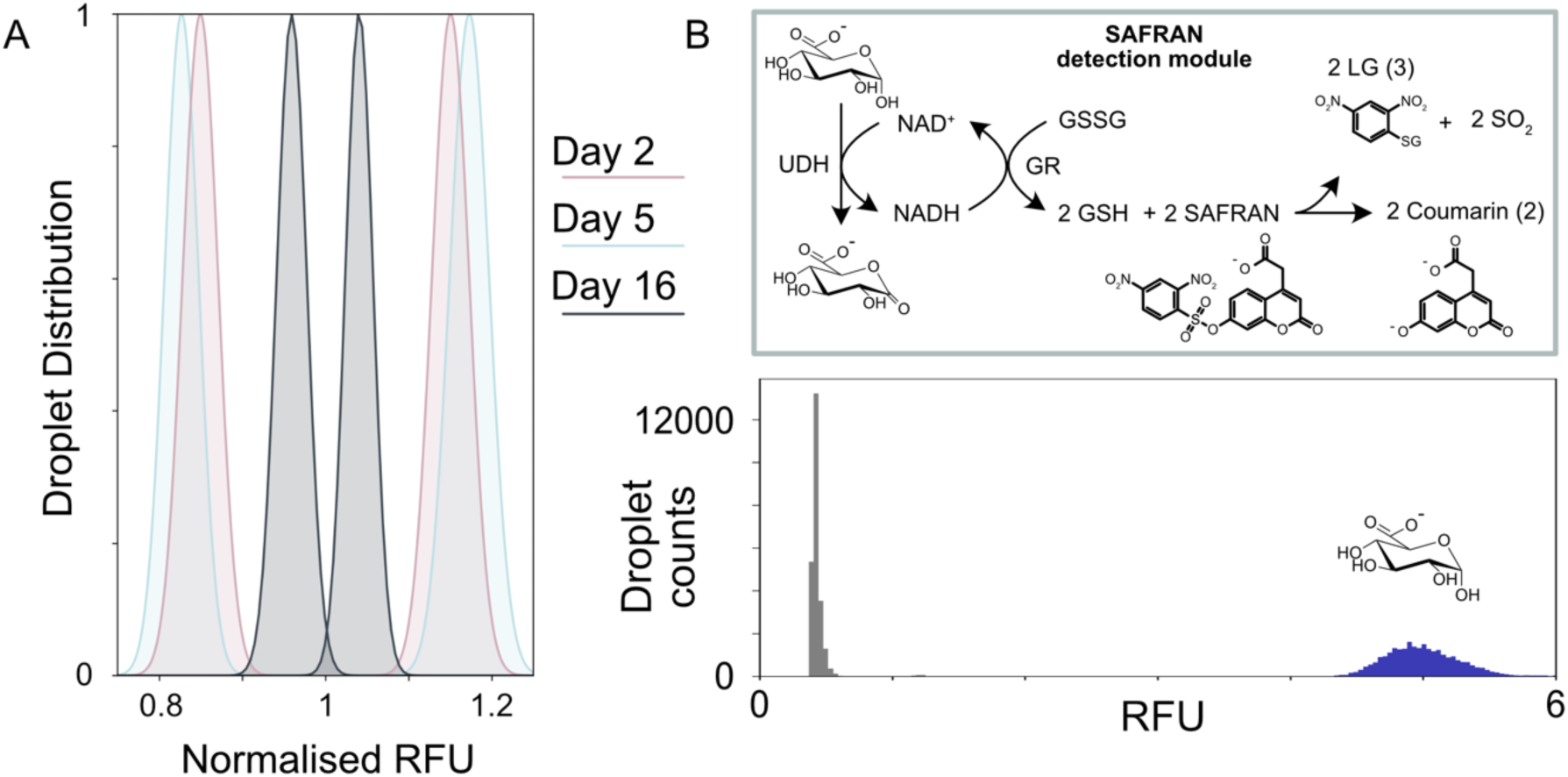
Droplet application of the coupled assay yields a stable signal that emerges rapidly. (A) Co-incubation of droplets containing 25 µM or 50 µM coumarin **2** for up to 16 days maintained two distinct fluorescent populations when measured on-chip. The data shown are the Gaussian fits to the mixed populations, when normalized by RFU and abundance for clarity of presentation. (B) Droplets with cascade reagents and 500 µM glucuronic acid or cascade reagents and buffer form two distinct populations on a FADS sorter when mixed, incubated for 40 min, and measured together. Despite the high analyte concentration, no droplet-to-droplet exchange of the product (or intermediates) of the reaction cascade that would jeopardise the distinction between positives and negatives is observed.

We then set out to test if oxidoreductase activity could be detected in droplets using SAFRAN by again creating a mixed two populations of droplets: one containing the SAFRAN cascade reagents, the oxidoreductase uronate dehydrogenase and glucuronic acid; the other identical but without glucuronic acid. Fluorescence was present at the limit of detection 10 minutes after droplet formation (**Figure S9**) but was distinct from background within 40 minutes (**Figure 3B**). Furthermore, these populations remained distinct after overnight incubation (**Figure S9**). These results demonstrate that the SAFRAN cascade can be used to detect the purified oxidoreductase uronate dehydrogenase in droplets, forming the basis for using these enzymes as coupling agents to detect upstream chemical reactions, such as the release of monosaccharides from polysaccharides.

Sensitivity is a major limitation of several droplet-based detection modes as for example absorbance-based detection (∼10 µM)^20,71^, which currently is the only method available for screening dehydrogenase activity using non-model substrates at ultrahigh throughput that does not suffer from leakage^59,70^. Fluorescent assays can be up to three orders of magnitude more sensitive, as shown by Colin *et al.* for a fluorescein-based phosphotriesterase assay with a detection limit of 2.5 nM^4^. This sensitivity allows detection of enzymatic activities with less than one turnover per enzyme molecule, thus enabling ultrahigh throughput screening even when expression or intrinsic activity are low. To test the limit of detection of the SAFRAN system using FADS, we sequentially measured the signal from droplets containing different concentrations of coumarin product **2** and found a linear dependence of fluorescence and product concentration with clear signal detectable down to 30 nM (corresponding to 542,000 molecules per droplet, or 271,000 turnovers of glutathione reductase, **Figure S10**). The lowest quantified droplet populations tested yielding distinct populations were 0 nM and 30 nM. (**Figure S10**). This sensitivity allows detection of enzymatic activities that lead to less than one turnover per enzyme molecule on average (considering that a single cell compartmentalised in a droplet typically produces µM enzyme^20^).

### Screening plasmid libraries of hydrolases and oxidoreductases against non-model substrates using FADS

Following confirmation that the SAFRAN system was amenable to extended droplet incubation, we investigated use of the system for screening enzyme libraries at ultrahigh throughput in an *E. coli* host. First, we tested the capacity of the system to identify glycosidases releasing monosaccharides, which served as the oxidisable substrate that was coupled to the SAFRAN cascade using a monosaccharide dehydrogenase. The two-step microfluidic workflow tested here is outlined in **Figure 4A**. This workflow requires expression of the cytoplasmic enzyme library in bulk media followed by co-encapsulation of cells with substrate, coupled assay and lysis reagents in a flow-focusing device. The droplets were then subjected to an incubation over a suitable timeframe to reach the limit of detection (at least 271,000 enzyme turnovers), followed by sorting using FADS. To test this workflow, a defined library consisting of a 1000:1 abundance of inactive GH115 E176A (the relative activities of the WT and E176A are shown in **Figure S11**) against a minority of wild-type enzyme, was screened for activity against a natural substrate, beechwood xylan (420 μM 4OMe-GlcA motif), as shown in **Figure 4B**. This library was encapsulated using a flow-focusing device and droplets were stored off-chip for 60 minutes, after which the most fluorescent 0.02% of droplets were sorted at 800 Hz using FADS **(Figure 4C and S12)**, yielding an enrichment of the library 765- fold, calculated according to the method of Baret *et al*.^72^ (**Figure S13**, **Supplementary Note 1**).

**Figure 4:**
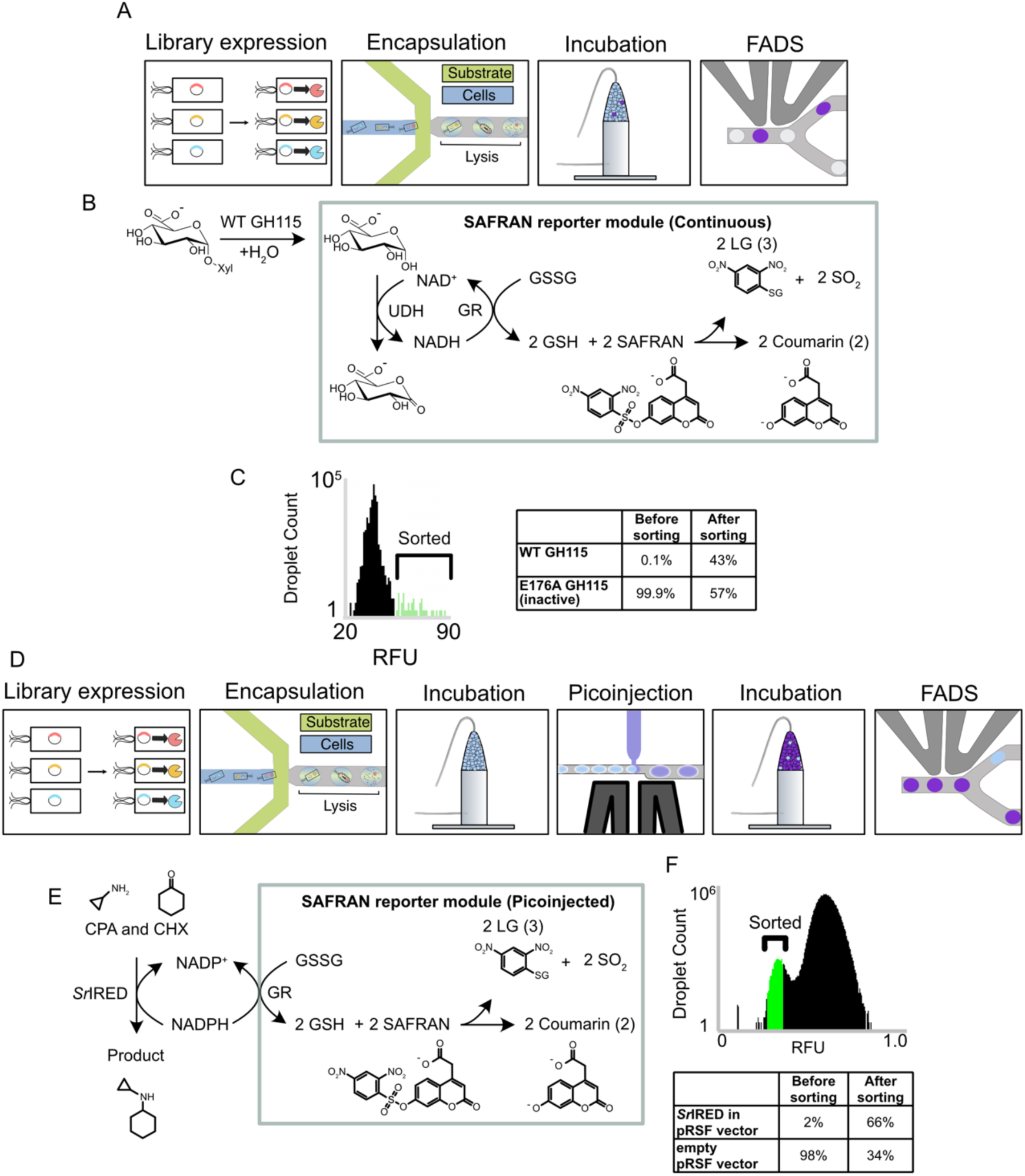
A sensitive droplet assay detects glycosidase and imine reductase activity from single cells on natural substrates. (A) Workflow used for testing the enrichment of glycosidases in droplets. (B) Schematic of the target droplet reaction in this assay. (C) (*Left*) Width-gated histogram describing a sample of droplet RFUs in the unsorted library. The top 0.02% of droplets were selected for sorting. (Right) Plate activity of sorted clones picked from the recovered library relative to the wild type. 39 of the 90 clones assayed were found to be confidently positive. (D) Workflow used for testing the enrichment of imine reductases in droplets. (E) Schematic of the target reaction in droplets. The imine reductase (IRED) first consumes NADPH until the droplets are picoinjected with the SAFRAN reporter module, which quenches the reaction and converts the unreacted NADPH to a fluorescent signal. (F) Droplet trace and enrichment statistics of the enrichment the IRED. Post-sorting library composition was determined using Sanger sequencing from twelve recovered clones.

To demonstrate quantification of the backwards reaction in droplets, where NADPH is consumed, the SAFRAN assay was used to enrich for cells expressing the bacterial imine reductase *Sr*IRED catalysing the condensation of CPA and CHX using the workflow outlined in **Figure 4D**. Individual bacteria containing either empty plasmid or plasmid containing the IRED gene where allowed to express the IRED, where present, in bulk media before mixing in a 98:2 ratio and co-encapsulation together with substrates and lysis reagents using a flow-focusing chip. After subsequent 1 hour off-chip incubation time, the droplets were picoinjected with the SAFRAN cascade components, which reported on the amount of unreacted NADPH remaining in each droplet (**Figure 4E**). Sorting with FADS for the low fluorescence population enabled an 98-fold enrichment of IREDs (**Figure 4F**), according to the method of Baret *et al*.^72^ (**Supplementary Note 2**).

The successful enrichments of enzymes from the hydrolase and the oxidoreductase reaction classes, using either the cofactor NADH or NADPH, measuring either cofactor production or consumption, establishes SAFRAN as a useful method for assaying any reaction that can be coupled to NAD(P)H consumption or production with extremely high sensitivity and at ultra-high throughput.

### Ultra-high throughput directed evolution of GH115 against the feedstock beechwood xylan

The SAFRAN assay delivers a promising system for performing directed enzyme evolution against a substrate of choice. The laboratory evolution of GH115 for higher activity against its substrate beechwood xylan promises direct applications in bioreactors for xylan utilization. In particular, improving kinetic parameters of the enzyme permit lower enzyme loadings and more efficient operation in complex mixtures of substrate. The enzyme used as a starting point, *AxyAgu*115A, is known to be a more tolerant enzyme to xylan substitutions than some other GH115 enzymes^73^, and possess some alkaline tolerance^33^. Alkaline conditions are preferable in xylan processing, as the solubility of the polymer is greater, but making enzymes compatible with these conditions remains a challenge^33^. Therefore, *AxyAgu*115A serves as a promising starting point for directed evolution of a more efficient GH115 enzyme.

For several reasons the GH115 family is a potentially challenging target for directed evolution: the enzymes are dimers of around 1000 amino acids per monomer and consist of 4-5 domains, with the second domain ‘B’ acting as the catalytic domain^73^. Although there is evidence that the enzyme acts *via* a Koshland inverting mechanism^74^, the catalytic acid and base have not been unambiguously identified. This is partially because the active sites are - unusual for a glycosidase - comprised of flexible loops, hindering crystallographic identification of the catalytic residues. The other domains are of unknown function, but are likely catalytically relevant as the active site alone domain is not sufficient for catalysis^75^.

We hypothesised that mutagenesis focused on domain B could target catalytic improvements, while the other four domains were left constant to retain protein stability and any other important yet cryptic roles. Whole-domain mutagenesis was achieved using error-prone PCR (**Figure 5A**). The substrate concentration in droplets was 420 µM of the (4-OMe)GlcA epitope, almost an order of magnitude below the *K*_M_ of the enzyme (1.49 mM), thus enacting selection pressure for variants that are able to strongly bind the substrate, targeting improved performance under bioreactor conditions were the epitope only comprises a small fraction of the total target plant cell wall loading. Following off-chip incubation of the library members and xylan for an hour, 3.6 million droplets were analysed, and the highest fluorescent droplets were sorted (**Figure 5B**). 39 wells from the output were identified as having higher than wild-type activity in cell lysate (43 %, **Figure 5C**), displaying up to 2-fold improvements in maximal release rate of (4-OMe)GlcA. Five members A-E were selected from the secondary screen for further characterization.

**Figure 5:**
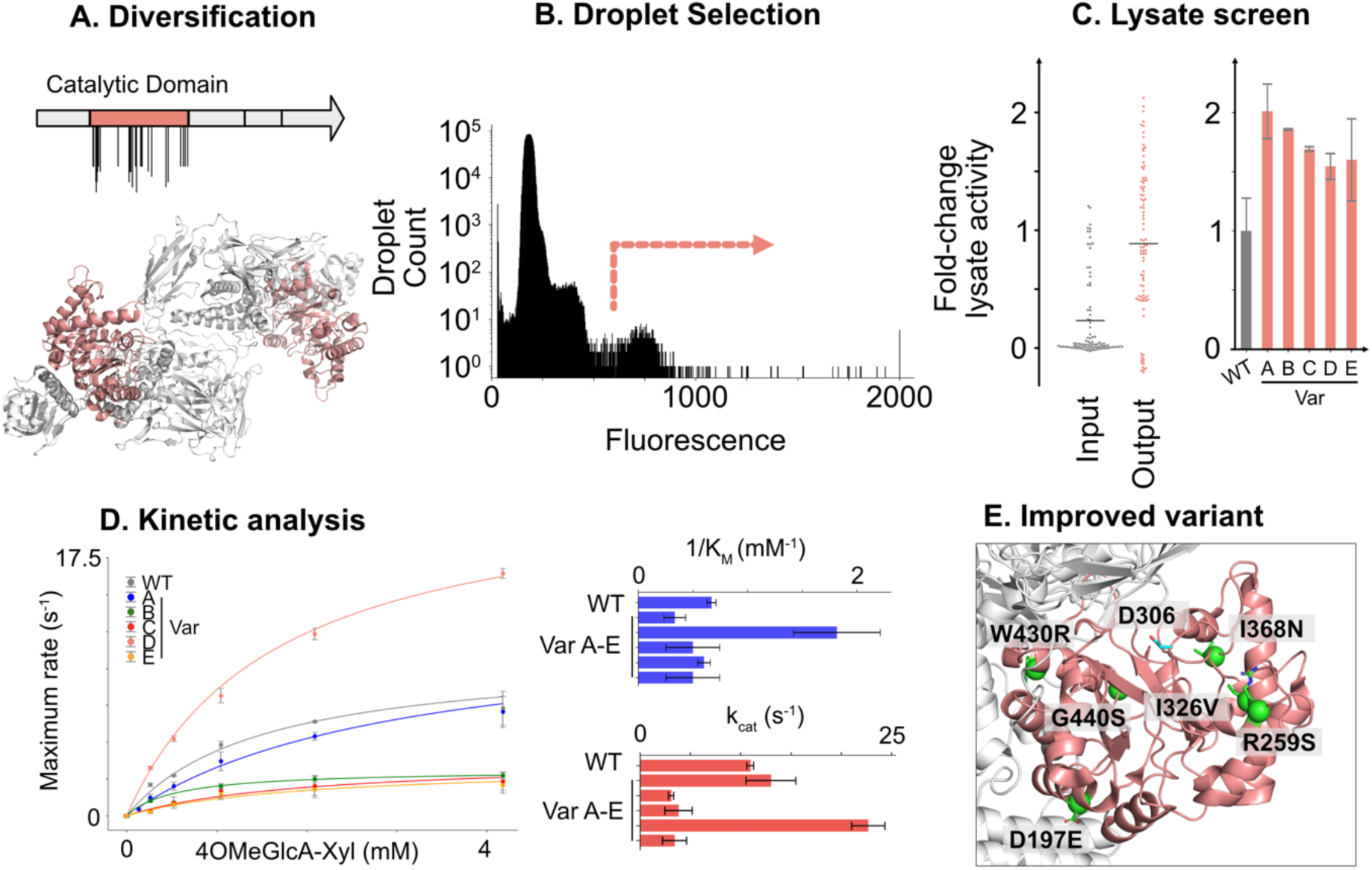
Directed evolution of GH115. (A) Schematic showing the dimeric structure of *AxyAgu115A* (PDB 6NPS), highlighting the catalytic domain. (B) Droplet trace from sort of 3.6 million droplet events. (C) (Left) Maximal rate of release of 4-OMeGlcA from BWX from sorted lysate from either the unsorted or sorted library in microtiter plates. Each point has been normalized between WT (=1) and inactive E176A (=0). (Right) Maximal rate of release of 4-OMeGlcA from BWX of the selected improving members. Values represent duplicate growth and measurement, except for wild type, which has six replicate growths and measurements. (D) Kinetic analysis of mutants identified as improving in lysate. Measurements are the average of three replicates. (E) Schematic showing the position of mutations in variant D (green) relative a putative catalytic residue (D306, cyan). Structure derives from PDB 6NPS.

These mutants were found to harbour up to six mutations (**Table 1**) and were isolated for further characterisation. L426P showed an almost 3-fold reduction in the K_M_ of the enzyme, while the mutant D197E R259S I326V I368N W430R G440S showed a 2.1-fold increase in k_cat_ and a non-significantly changed K_M_, resulting in 1.9-fold increase in k_cat_/K_M_ (**Table 1**, **Figure 5D**). This variant contains mutations scattered around the catalytic domain, with position R259 notably interacting with the disordered loop shown to be critical for catalysis (**Figure 5E**). By circumventing the dependence on modified substrates, this single round of directed evolution *via* SAFRAN coupled assay achieves amongst the highest reported improvements in kinetic parameters of glycosidases against their unmodified substrates (**Table S1**).

**Table 1:**
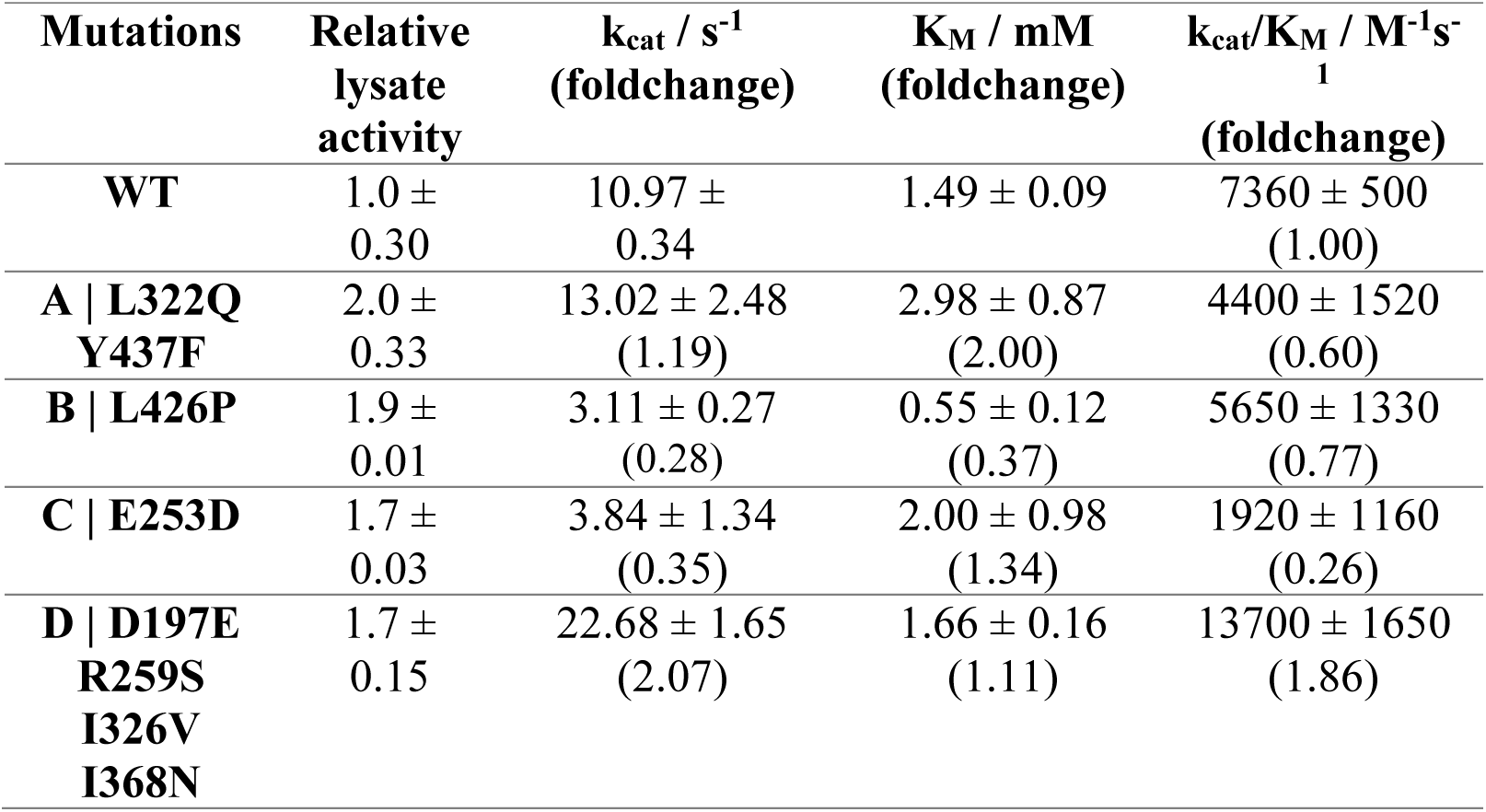
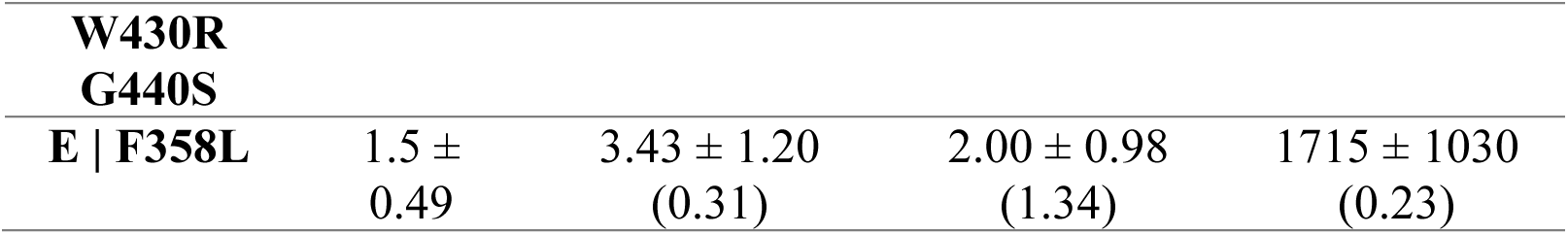
Summary of purified enzyme parameters derived from five selected output members.

## Discussion

Our ultra-high throughput fluorogenic NAD(P)H assay enables large-scale screening of enzyme libraries against an unmodified substrate of interest, which we have applied here for enzyme engineering by directed evolution. We achieved this by designing a caged coumarin, SAFRAN, that is uncaged following NAD(P)H-dependant reduction of oxidised glutathione, and subsequent reaction with the free thiols of reduced glutathione. The coumarin product carries a carboxyl group, preventing leakage between droplets. We demonstrate that screening in an *E. coli* expression host is compatible with this cascade, with the assay signal overcoming any interference derived from endogenous *E. coli* reduced glutathione and NAD(P)H reserves.

We use the cascade to meet the challenge of engineering glycosyl hydrolases, which alongside medical applications^76^ are an important class of enzymes for their ability to convert biomass into valuable small molecules with high specificity and selectivity^77^. The enzyme *AxyAgu*115A has recently been used in a cascade reaction to convert hardwood glucuronoxylan, an underused fraction from biorefineries, to the dicarboxylic acid 4-O-methyl D-glucaric acid^78^. Glucuronic acid and its derivatives are valuable chemicals for synthetic purposes and demand currently for these compounds far outstrips the current capacities of chemical synthesis and extraction from natural sources^79^. Glucuronic acid is also being used to phase out less sustainable chemicals in manufacturing; dicarboxylic acids are valuable building blocks for synthesis of pharmaceuticals and bioplastics, but demand is currently mostly met using peterochemcials^80^. The approach described in this work enabled us to select *AxyAgu*115A variants for their activity against their natural substrate – the feedstock glucuronoxylan – and not against an activated model substrate, thus ensuring that improvements made during the directed evolution translate to industrially relevant catalytic improvements against the polymer of interest.

The application of this assay to insoluble substrates in microfluidic droplets – common targets of interest for industrial applications^81^ – remains to be demonstrated. Unlike the soluble substrates tested in this work, insoluble particles pose challenges of (i) sedimentation (ii) optical interference with fluorophores and (iii) destabilization of the oil-aqueous interface. One solution already in use to overcome these challenges are microscale hydrogel beads^82^. Hydrogels immobilize particles such as insoluble substrates and even whole cells^83^, and are compatible with fluorescence detection methods^84^. The immobilization of substrate in the hydrogel stabilizes the surrounding droplet-oil interface. Hydrogel formats therefore pose an attractive avenue to expand the SAFRAN coupled assay to screening insoluble substrates.

The modest rate accelerations achieved by directed evolution have to be evaluated in the context of the engineering challenges in the GH115 family. Family members are large dimers, with each monomer containing up to 1000 amino acids. Furthermore, beyond dimerization, the functions of the domains other than the TIM-barrel catalytic domain have yet to be elucidated. Despite the family seemingly operating an inverting Koshland mechanism^74^, the active site residues have also not been decisively identified, and are likely present on the long and flexible loops that cover the active site and have been implicated in catalysis and substrate scope^73^. Using the available information on the GH115 family we focused mutagenesis on the active site domain and found that the most improving variant had mutations not located in the active site, but in the second shell residues and in more distant positions throughout the catalytic domain. The position R259, mutated to serine in the improved mutant, is a second-shell residue known to form a salt bridge with the long active site loop and thought to influence the tolerance of the enzyme to substitutions on the xylan backbone^73^. W430 (mutated to arginine) is found at the domain interface of the catalytic domain and two other domains within the protein, potentially influencing long-range protein dynamics.

While we demonstrate the cascade for engineering *AxyAgu*115A, the development of the SAFRAN cascade was undertaken with the primary goal of making dehydrogenase coupled assays amenable to droplet screening. Based on the assay’s modularity, we propose that other dehydrogenase-coupled assays can now be analysed using FADS (or, after a second emulsification step, by standard flow cytometric sorting in double emulsions^85,86^), taking advantage of the wealth of literature that reports on coupling reactions to NAD(P)H production or depletion. These include xylosidases, glucosidases, arabinosidases^70^, small-molecule/natural product methyltransferases^87^, nucleotide methyltransferases^88^, ATPases and other enzymes that produce ADP^89,90^ including kinases^91^, systems that produce urea or pyruvate^92^, amino acid oxidative deaminases^93^, phosphite dehydrogenases^94^ and alkane hydroxylases^95^. Additionally, engineering of redox enzymes that are not in a coupled set-up, such dehydrogenases for biocatalytic applications^96,97^, including conversion of monosaccharides to hydrogen gas^98^, will directly benefit from this sensitive screening assay. As a proof-of-principle we demonstrate how the SAFRAN assay is suitable for screening libraries of imine reductases, a pharmaceutically valuable synthetic family of enzymes.

Based on these degrees of freedom, SAFRAN is a uniquely modular detection system, where the same assay reagent can be deployed in various contexts, the elements of which can be modularly exchanged: (i) On the one hand, a wide range of primary substrates can be detected, as long as they lead to an intermediate that is processed by the coupled reaction. For example, reactions of a wide range of natural glycosides can be detected, as long monosaccharides are generated that feed into a coupling reaction. Therefore, the range of options for substrate detection is determined by the availability of specific *upstream* coupling enzymes that can be modularly exchanged. (ii) On the other hand, the reaction type can be chosen by employing any of the abovementioned enzymes that consume or generates NAD(P)H, here by exchanging the *downstream* coupling enzymes that process the intermediate substrate in redox reaction.

## Conclusion

In contrast to alternative NADPH detection strategies, the improved sensitivity of our approach^20^ and reduced leakage between droplets^30^ allows the detection of low-activity enzymes by extended droplet incubation times. This is crucial for protein engineering and discovery of other biocatalysts^99–101^, as the recruitment of proteins for the conversion of our current economic model to a sustainable bioeconomy depends on three sources with typically weak activities. These are (i) promiscuous side activities of existing enzymes^102,103^, (ii) enzymes from metagenomic sources^104^, identified by functional screening or by bioinformatics and (iii) computationally designed enzymes^99–101^. Without a sensitive assay, these useful, but imperfect starting points cannot be improved by directed evolution and compatibility with an ultrahigh throughput method is crucial for success. Droplet microfluidics is arguably one of the most powerful UHT formats, allowing screening of ∼10^7^-membered libraries in a day, and SAFRAN makes sensitive fluorogenic screening of a broad range of reactions possible in this format.

## Acknowledgements

The authors thank Prof. Emma Master for a gene string for *AxyAgu*115A, Dr Tomasz Kaminski for microfluidic chip fabrication and members of the Hollfelder lab for helpful discussions and comments. This work was supported by the European Union’s Horizon 2020 research and innovation programme *via* an ERC Advanced Investigator grant (to FH; 695669), by the Biological and Biotechnological Research Council (BB/W006391/1), the Volkswagen Foundation (98182) and the Horizon Europe projects BlueTools (101081957) and BlueRemediomics (101082304). M.P was supported by a BBSRC iCASE studentship supported by Novozymes (BBSRC BB/X010899/1). O.J.K. was supported by an EPSRC studentship (EP/R513180/1). F.E.H.N. was supported by the European Union Marie-Curie network ‘Oligomed’ (956070). S.B. and P.B. thank the Chemistry Department at Cambridge University for financial support.

## Notes

### Competing Interest Statement

The authors have declared no competing interest.

### Summary of Updates

Additional experiments to show versatility of the detection system in both directions of cofactor redox reaction.

## References

(1) Arnold, F. H. Innovation by Evolution: Bringing New Chemistry to Life (Nobel Lecture). Angew Chem Int Ed 2019, 58 (41), 14420–14426. 10.1002/anie.201907729.

(2) Radley, E.; Davidson, J.; Foster, J.; Obexer, R.; Bell, E. L.; Green, A. P. Engineering Enzymes for Environmental Sustainability. Angew. Chem. Int. Ed. n/a (n/a), e202309305. 10.1002/anie.202309305.

(3) Lorenz, P.; Eck, J. Metagenomics and Industrial Applications. Nat. Rev. Microbiol. 2005, 3 (6), 510–516. 10.1038/nrmicro1161.

(4) Colin, P.-Y.; Kintses, B.; Gielen, F.; Miton, C. M.; Fischer, G.; Mohamed, M. F.; Hyvönen, M.; Morgavi, D. P.; Janssen, D. B.; Hollfelder, F. Ultrahigh-Throughput Discovery of Promiscuous Enzymes by Picodroplet Functional Metagenomics. Nat. Commun. 2015, 6 (1), 10008. 10.1038/ncomms10008.

(5) Gantz, M.; Aleku, G. A.; Hollfelder, F. Ultrahigh-Throughput Screening in Microfluidic Droplets: A Faster Route to New Enzymes. Trends Biochem. Sci. 2022, 47 (5), 451–452. 10.1016/j.tibs.2021.11.001.

(6) Gantz, M.; Neun, S.; Medcalf, E. J.; van Vliet, L. D.; Hollfelder, F. Ultrahigh-Throughput Enzyme Engineering and Discovery in In Vitro Compartments. Chem. Rev. 2023, 123 (9), 5571–5611. 10.1021/acs.chemrev.2c00910.

(7) Neun, S.; Brear, P.; Campbell, E.; Tryfona, T.; El Omari, K.; Wagner, A.; Dupree, P.; Hyvönen, M.; Hollfelder, F. Functional Metagenomic Screening Identifies an Unexpected β-Glucuronidase. Nat. Chem. Biol. 2022, 18 (10), 1096–1103. 10.1038/s41589-022-01071-x.

(8) Schnettler, J. D.; Klein, O. J.; Kaminski, T. S.; Colin, P.-Y.; Hollfelder, F. Ultrahigh-Throughput Directed Evolution of a Metal-Free α/β-Hydrolase with a Cys-His-Asp Triad into an Efficient Phosphotriesterase. J. Am. Chem. Soc. 2023, 145 (2), 1083–1096. 10.1021/jacs.2c10673.

(9) Schnettler, J. D.; Wang, M. S.; Gantz, M.; Karas, C.; Hollfelder, F.; Hecht, M. H. Selection of a Promiscuous Minimalist cAMP Phosphodiesterase from a Library of De Novo Designed Proteins. bioRxiv February 16, 2023, p 2023.02.13.528392. 10.1101/2023.02.13.528392.

(10) Obexer, R.; Godina, A.; Garrabou, X.; Mittl, P. R. E.; Baker, D.; Griffiths, A. D.; Hilvert, D. Emergence of a Catalytic Tetrad during Evolution of a Highly Active Artificial Aldolase. Nat. Chem. 2017, 9 (1), 50–56. 10.1038/nchem.2596.

(11) Ma, F.; Chung, M. T.; Yao, Y.; Nidetz, R.; Lee, L. M.; Liu, A. P.; Feng, Y.; Kurabayashi, K.; Yang, G.-Y. Efficient Molecular Evolution to Generate Enantioselective Enzymes Using a Dual-Channel Microfluidic Droplet Screening Platform. Nat. Commun. 2018, 9 (1), 1030. 10.1038/s41467-018-03492-6.

(12) Chen, H.-M.; Nasseri, S. A.; Rahfeld, P.; Wardman, J. F.; Kohsiek, M.; Withers, S. G. Synthesis and Evaluation of Sensitive Coumarin-Based Fluorogenic Substrates for Discovery of α-N-Acetyl Galactosaminidases through Droplet-Based Screening. Org. Biomol. Chem. 2021, 19 (4), 789–793. 10.1039/D0OB02484H.

(13) You, L.; Arnold, F. H. Directed Evolution of Subtilisin E in Bacillus Subtilis to Enhance Total Activity in Aqueous Dimethylformamide. Protein Eng. 1996, 9 (1), 77–83. 10.1093/protein/9.1.77.

(14) Wang, M.; Si, T.; Zhao, H. Biocatalyst Development by Directed Evolution. Bioresour. Technol. 2012, *115C*, 117–125. 10.1016/j.biortech.2012.01.054.

(15) Hecko, S.; Schiefer, A.; Badenhorst, C. P. S.; Fink, M. J.; Mihovilovic, M. D.; Bornscheuer, U. T.; Rudroff, F. Enlightening the Path to Protein Engineering: Chemoselective Turn-On Probes for High-Throughput Screening of Enzymatic Activity. Chem. Rev. 2023, 123 (6), 2832–2901. 10.1021/acs.chemrev.2c00304.

(16) Menke, M. J.; Pascal Schneider, P.; Badenhorst, C. P. S.; Kunzendorf, A.; Heinz, F.; Dörr, M.; Hayes, M. A.; Bornscheuer, U. A Universal, Continuous Assay for SAM-Dependent Methyltransferases. Angew. Chem. Int. Ed. n/a (n/a), e202313912. 10.1002/anie.202313912.

(17) Bauer, J. A.; Zámocká, M.; Majtán, J.; Bauerová-Hlinková, V. Glucose Oxidase, an Enzyme “Ferrari”: Its Structure, Function, Production and Properties in the Light of Various Industrial and Biotechnological Applications. Biomolecules 2022, 12 (3), 472. 10.3390/biom12030472.

(18) Garcia-Hernandez, C.; Garcia-Cabezon, C.; Martin-Pedrosa, F.; Rodriguez-Mendez, M. L. Analysis of Musts and Wines by Means of a Bio-Electronic Tongue Based on Tyrosinase and Glucose Oxidase Using Polypyrrole/Gold Nanoparticles as the Electron Mediator. Food Chem. 2019, 289, 751–756. 10.1016/j.foodchem.2019.03.107.

(19) Megazyme. D-glucuronic acid & d-galacturonic acid (d-glucuronate & d-galacturonate) assay procedure. https://www.megazyme.com/documents/Assay_Protocol/K-URONIC_DATA.pdf (accessed 2023-09-01).

(20) Gielen, F.; Hours, R.; Emond, S.; Fischlechner, M.; Schell, U.; Hollfelder, F. Ultrahigh-Throughput-Directed Enzyme Evolution by Absorbance-Activated Droplet Sorting (AADS). Proc. Natl. Acad. Sci. U. S. A. 2016, 113 (47), E7383–E7389. 10.1073/pnas.1606927113.

(21) Marshall, J. R.; Yao, P.; Montgomery, S. L.; Finnigan, J. D.; Thorpe, T. W.; Palmer, R. B.; Mangas-Sanchez, J.; Duncan, R. A. M.; Heath, R. S.; Graham, K. M.; Cook, D. J.; Charnock, S. J.; Turner, N. J. Screening and Characterization of a Diverse Panel of Metagenomic Imine Reductases for Biocatalytic Reductive Amination. Nat. Chem. 2021, 13 (2), 140–148. 10.1038/s41557-020-00606-w.

(22) Zurek, P. J.; Knyphausen, P.; Neufeld, K.; Pushpanath, A.; Hollfelder, F. UMI-Linked Consensus Sequencing Enables Phylogenetic Analysis of Directed Evolution. Nat. Commun. 2020, 11 (1), 6023. 10.1038/s41467-020-19687-9.

(23) Principles of Fluorescence Spectroscopy, 3rd ed.; Lakowicz, J. R., Ed.; Springer US: Boston, MA, 2006. 10.1007/978-0-387-46312-4.

(24) Gorbunova, I. A.; Danilova, M. K.; Sasin, M. E.; Belik, V. P.; Golyshev, D. P.; Vasyutinskii, O. S. Determination of Fluorescence Quantum Yields and Decay Times of NADH and FAD in Water–Alcohol Mixtures: The Analysis of Radiative and Nonradiative Relaxation Pathways. J. Photochem. Photobiol. Chem. 2023, 436, 114388. 10.1016/j.jphotochem.2022.114388.

(25) Swoboda, A.; Pfeifenberger, L. J.; Duhović, Z.; Bürgler, M.; Oroz-Guinea, I.; Bangert, K.; Weißensteiner, F.; Parigger, L.; Ebner, K.; Glieder, A.; Kroutil, W. Enantioselective High-Throughput Assay Showcased for the Identification of (R)- as Well as (S)-Selective Unspecific Peroxygenases for C-H Oxidation. Angew. Chem. Int. Ed Engl. 2023, 62 (46), e202312721. 10.1002/anie.202312721.

(26) Velick, S. F. Fluorescence Spectra and Polarization of Glyceraldehyde-3-Phosphate and Lactic Dehydrogenase Coenzyme Complexes. J. Biol. Chem. 1958, 233 (6), 1455–1467. 10.1016/S0021-9258(18)49355-6.

(27) Brochon, J. C.; Wahl, P.; Monneuse-Doublet, M. O.; Olomucki, A. Pulse Fluorimetry Study of Octopine Dehydrogenase-Reduced Nicotinamide Adenine Dinucleotide Complexes. Biochemistry 1977, 16 (21), 4594–4599. 10.1021/bi00640a010.

(28) Cannon, T. M.; Lagarto, J. L.; Dyer, B. T.; Garcia, E.; Kelly, D. J.; Peters, N. S.; Lyon, A. R.; French, P. M. W.; Dunsby, C. Characterization of NADH Fluorescence Properties under One-Photon Excitation with Respect to Temperature, pH, and Binding to Lactate Dehydrogenase. Osa Contin. 2021, 4 (5), 1610–1625. 10.1364/OSAC.423082.

(29) Ma, N.; Digman, M. A.; Malacrida, L.; Gratton, E. Measurements of Absolute Concentrations of NADH in Cells Using the Phasor FLIM Method. Biomed. Opt. Express 2016, 7 (7), 2441–2452. 10.1364/BOE.7.002441.

(30) Scheler, O.; Kaminski, T. S.; Ruszczak, A.; Garstecki, P. Dodecylresorufin (C12R) Outperforms Resorufin in Microdroplet Bacterial Assays. ACS Appl. Mater. Interfaces 2016, 8 (18), 11318–11325. 10.1021/acsami.6b02360.

(31) Ryabova, O.; Vrsanská, M.; Kaneko, S.; van Zyl, W. H.; Biely, P. A Novel Family of Hemicellulolytic Alpha-Glucuronidase. FEBS Lett. 2009, 583 (9), 1457–1462. 10.1016/j.febslet.2009.03.057.

(32) Mortimer, J. C.; Miles, G. P.; Brown, D. M.; Zhang, Z.; Segura, M. P.; Weimar, T.; Yu, X.; Seffen, K. A.; Stephens, E.; Turner, S. R.; Dupree, P. Absence of Branches from Xylan in Arabidopsis Gux Mutants Reveals Potential for Simplification of Lignocellulosic Biomass. Proc. Natl. Acad. Sci. 2010, 107 (40), 17409–17414. 10.1073/pnas.1005456107.

(33) Yan, R.; Vuong, T. V.; Wang, W.; Master, E. R. Action of a GH115 α-Glucuronidase from Amphibacillus Xylanus at Alkaline Condition Promotes Release of 4-O- Methylglucopyranosyluronic Acid from Glucuronoxylan and Arabinoglucuronoxylan. Enzyme Microb. Technol. 2017, 104, 22–28. 10.1016/j.enzmictec.2017.05.004.

(34) Ravn, J. L.; Manfrão-Netto, J. H. C.; Schaubeder, J. B.; Torello Pianale, L.; Spirk, S.; Ciklic, I. F.; Geijer, C. Engineering Saccharomyces Cerevisiae for Targeted Hydrolysis and Fermentation of Glucuronoxylan through CRISPR/Cas9 Genome Editing. Microb. Cell Factories 2024, 23 (1), 85. 10.1186/s12934-024-02361-w.

(35) Raji, O.; Arnling Bååth, J.; Vuong, T. V.; Larsbrink, J.; Olsson, L.; Master, E. R. The Coordinated Action of Glucuronoyl Esterase and α-Glucuronidase Promotes the Disassembly of Lignin–Carbohydrate Complexes. FEBS Lett. 2021, 595 (3), 351–359. 10.1002/1873-3468.14019.

(36) Banerjee, G.; Car, S.; Scott-Craig, J. S.; Borrusch, M. S.; Aslam, N.; Walton, J. D. Synthetic Enzyme Mixtures for Biomass Deconstruction: Production and Optimization of a Core Set. Biotechnol. Bioeng. 2010, 106 (5), 707–720. 10.1002/bit.22741.

(37) Perna, V. N.; Barrett, K.; Meyer, A. S.; Zeuner, B. Substrate Specificity and Transglycosylation Capacity of α-L-Fucosidases across GH29 Assessed by Bioinformatics-Assisted Selection of Functional Diversity. Glycobiology 2023, 33 (5), 396–410. 10.1093/glycob/cwad029.

(38) Martínez Gascueña, A.; Wu, H.; Wang, R.; Owen, C. D.; Hernando, P. J.; Monaco, S.; Penner, M.; Xing, K.; Le Gall, G.; Gardner, R.; Ndeh, D.; Urbanowicz, P. A.; Spencer, D. I. R.; Walsh, M.; Angulo, J.; Juge, N. Exploring the Sequence-Function Space of Microbial Fucosidases. Commun. Chem. 2024, 7 (1), 1–15. 10.1038/s42004-024-01212-4.

(39) Pidatala, V. R.; Mahboubi, A.; Mortimer, J. C. Structural Characterization of Mannan Cell Wall Polysaccharides in Plants Using PACE. J. Vis. Exp. JoVE 2017, No. 128, 56424. 10.3791/56424.

(40) Santos, C. A.; Morais, M. A. B.; Terrett, O. M.; Lyczakowski, J. J.; Zanphorlin, L. M.; Ferreira-Filho, J. A.; Tonoli, C. C. C.; Murakami, M. T.; Dupree, P.; Souza, A. P. An Engineered GH1 β-Glucosidase Displays Enhanced Glucose Tolerance and Increased Sugar Release from Lignocellulosic Materials. Sci. Rep. 2019, 9, 4903. 10.1038/s41598-019-41300-3.

(41) Chao, L.; Jongkees, S. High-Throughput Approaches in Carbohydrate-Active Enzymology: Glycosidase and Glycosyl Transferase Inhibitors, Evolution, and Discovery. Angew. Chem. Int. Ed. 2019, 58 (37), 12750–12760. 10.1002/anie.201900055.

(42) Sellés Vidal, L.; Kelly, C. L.; Mordaka, P. M.; Heap, J. T. Review of NAD(P)H-Dependent Oxidoreductases: Properties, Engineering and Application. Biochim. Biophys. Acta BBA - Proteins Proteomics 2018, 1866 (2), 327–347. 10.1016/j.bbapap.2017.11.005.

(43) The Year In New Drugs. Chemical & Engineering News. https://cen.acs.org/articles/94/i5/Year-New-Drugs.html (accessed 2024-08-03).

(44) Gantz, M.; Mathis, S. V.; Nintzel, F. E. H.; Zurek, P. J.; Knaus, T.; Patel, E.; Boros, D.; Weberling, F.-M.; Kenneth, M. R. A.; Klein, O. J.; Medcalf, E. J.; Moss, J.; Herger, M.; Kaminski, T. S.; Mutti, F. G.; Lio, P.; Hollfelder, F. Microdroplet Screening Rapidly Profiles a Biocatalyst to Enable Its AI-Assisted Engineering. bioRxiv April 8, 2024, p 2024.04.08.588565. 10.1101/2024.04.08.588565.

(45) Goubet, F.; Barton, C. J.; Mortimer, J. C.; Yu, X.; Zhang, Z.; Miles, G. P.; Richens, J.; Liepman, A. H.; Seffen, K.; Dupree, P. Cell Wall Glucomannan in Arabidopsis Is Synthesised by CSLA Glycosyltransferases, and Influences the Progression of Embryogenesis. Plant J. 2009, 60 (3), 527–538. 10.1111/j.1365-313X.2009.03977.x.

(46) Temple, H.; Phyo, P.; Yang, W.; Lyczakowski, J. J.; Echevarría-Poza, A.; Yakunin, I.; Parra-Rojas, J. P.; Terrett, O. M.; Saez-Aguayo, S.; Dupree, R.; Orellana, A.; Hong, M.; Dupree, P. Golgi-Localized Putative S-Adenosyl Methionine Transporters Required for Plant Cell Wall Polysaccharide Methylation. Nat. Plants 2022, 8 (6), 656–669. 10.1038/s41477-022-01156-4.

(47) Zhang, B.; Ge, C.; Yao, J.; Liu, Y.; Xie, H.; Fang, J. Selective Selenol Fluorescent Probes: Design, Synthesis, Structural Determinants, and Biological Applications. J. Am. Chem. Soc. 2015, 137 (2), 757–769. 10.1021/ja5099676.

(48) Shao, A.; Xu, Q.; Kang, C. W.; Cain, C. F.; Lee, A. C.; Tang, C.-H. A.; Del Valle, J. R.; Hu, C.-C. A. IRE-1-Targeting Caged Prodrug with Endoplasmic Reticulum Stress-Inducing and XBP-1S-Inhibiting Activities for Cancer Therapy. Mol. Pharm. 2022, 19 (4), 1059–1067. 10.1021/acs.molpharmaceut.1c00639.

(49) Maeda, H.; Matsuno, H.; Ushida, M.; Katayama, K.; Saeki, K.; Itoh, N. 2,4- Dinitrobenzenesulfonyl Fluoresceins as Fluorescent Alternatives to Ellman’s Reagent in Thiol-Quantification Enzyme Assays. Angew. Chem. Int. Ed. 2005, 44 (19), 2922–2925. 10.1002/anie.200500114.

(50) Roubinet, B.; Renard, P.-Y.; Romieu, A. New Insights into the Water-Solubilization of Thiol-Sensitive Fluorogenic Probes Based on Long-Wavelength 7-Hydroxycoumarin Scaffolds. Dyes Pigments 2014, 110, 270–284. 10.1016/j.dyepig.2014.02.004.

(51) Basavarajaiah, S. M.; Gunavanthrao Yernale, N.; Punith Kumar, M.; Rakesh, B. Review on Contemporary Synthetic Recipes to Access Versatile Coumarin Heterocycles. Polycycl. Aromat. Compd. 2023, 0 (0), 1–25. 10.1080/10406638.2023.2235874.

(52) Bouhaoui, A.; Eddahmi, M.; Dib, M.; Khouili, M.; Aires, A.; Catto, M.; Bouissane, L. Synthesis and Biological Properties of Coumarin Derivatives. A Review. ChemistrySelect 2021, 6 (24), 5848–5870. 10.1002/slct.202101346.

(53) Zinchenko, A.; Devenish, S. R. A.; Hollfelder, F. Rapid Quantitative Assessment of Small Molecule Leakage from Microdroplets by Flow Cytometry and Improvement of Fluorophore Retention in Biochemical Assays. bioRxiv April 23, 2023, p 2023.04.23.538007. 10.1101/2023.04.23.538007.

(54) Debon, A.; Pott, M.; Obexer, R.; Green, A. P.; Friedrich, L.; Griffiths, A. D.; Hilvert, D. Ultrahigh-Throughput Screening Enables Efficient Single-Round Oxidase Remodelling. Nat. Catal. 2019, 2 (9), 740–747. 10.1038/s41929-019-0340-5.

(55) Debon, A. P.; Wootton, R. C. R.; Elvira, K. S. Droplet Confinement and Leakage: Causes, Underlying Effects, and Amelioration Strategies. Biomicrofluidics 2015, 9 (2), 024119. 10.1063/1.4917343.

(56) Payne, E. M.; Taraji, M.; Murray, B. E.; Holland-Moritz, D. A.; Moore, J. C.; Haddad, P. R.; Kennedy, R. T. Evaluation of Analyte Transfer between Microfluidic Droplets by Mass Spectrometry. Anal. Chem. 2023, 95 (10), 4662–4670. 10.1021/acs.analchem.2c04985.

(57) Courtois, F.; Olguin, L. F.; Whyte, G.; Theberge, A. B.; Huck, W. T. S.; Hollfelder, F.; Abell, C. Controlling the Retention of Small Molecules in Emulsion Microdroplets for Use in Cell-Based Assays. Anal. Chem. 2009, 81 (8), 3008–3016. 10.1021/ac802658n.

(58) Gimeno-Pérez, M.; Finnigan, J. D.; Echeverria, C.; Charnock, S. J.; Hidalgo, A.; Mate, D. M. A Coupled Ketoreductase-Diaphorase Assay for the Detection of Polyethylene Terephthalate-Hydrolyzing Activity. ChemSusChem 2022, 15 (9), e202102750. 10.1002/cssc.202102750.

(59) Klaus, M.; Zurek, P. J.; Kaminski, T. S.; Pushpanath, A.; Neufeld, K.; Hollfelder, F. Ultrahigh-Throughput Detection of Enzymatic Alcohol Dehydrogenase Activity in Microfluidic Droplets with a Direct Fluorogenic Assay. ChemBioChem 2021, 22 (23), 3292–3299. 10.1002/cbic.202100322.

(60) Fink, D. W.; Koehler, W. R. pH Effects on Fluorescence of Umbelliferone. Anal. Chem. 1970, 42 (9), 990–993. 10.1021/ac60291a034.

(61) Scrutton, N. S.; Berry, A.; Perham, R. N. Redesign of the Coenzyme Specificity of a Dehydrogenase by Protein Engineering. Nature 1990, 343 (6253), 38–43. 10.1038/343038a0.

(62) Cataldi, T. R. I.; Nardiello, D. Determination of Free Proline and Monosaccharides in Wine Samples by High-Performance Anion-Exchange Chromatography with Pulsed Amperometric Detection (HPAEC-PAD). J. Agric. Food Chem. 2003, 51 (13), 3737–3742. 10.1021/jf034069c.

(63) Kurzyna-Szklarek, M.; Cybulska, J.; Zdunek, A. Analysis of the Chemical Composition of Natural Carbohydrates – An Overview of Methods. Food Chem. 2022, 394, 133466. 10.1016/j.foodchem.2022.133466.

(64) Deshavath, N. N.; Mukherjee, G.; Goud, V. V.; Veeranki, V. D.; Sastri, C. V. Pitfalls in the 3, 5-Dinitrosalicylic Acid (DNS) Assay for the Reducing Sugars: Interference of Furfural and 5-Hydroxymethylfurfural. Int. J. Biol. Macromol. 2020, 156, 180–185. 10.1016/j.ijbiomac.2020.04.045.

(65) Geiss, A. F.; Reichhart, T. M. B.; Pejker, B.; Plattner, E.; Herzog, P. L.; Schulz, C.; Ludwig, R.; Felice, A. K. G.; Haltrich, D. Engineering the Turnover Stability of Cellobiose Dehydrogenase toward Long-Term Bioelectronic Applications. ACS Sustain. Chem. Eng. 2021, 9 (20), 7086–7100. 10.1021/acssuschemeng.1c01165.

(66) Zhang, Z.; Khan, N. M.; Nunez, K. M.; Chess, E. K.; Szabo, C. M. Complete Monosaccharide Analysis by High-Performance Anion-Exchange Chromatography with Pulsed Amperometric Detection. Anal. Chem. 2012, 84 (9), 4104–4110. 10.1021/ac300176z.

(67) Macdonald, S. S.; Armstrong, Z.; Morgan-Lang, C.; Osowiecka, M.; Robinson, K.; Hallam, S. J.; Withers, S. G. Development and Application of a High-Throughput Functional Metagenomic Screen for Glycoside Phosphorylases. Cell Chem. Biol. 2019, 26 (7), 1001–1012.e5. 10.1016/j.chembiol.2019.03.017.

(68) Baret, J.-C. Surfactants in Droplet-Based Microfluidics. Lab. Chip 2012, 12 (3), 422–433. 10.1039/C1LC20582J.

(69) Sela, Y.; Magdassi, S.; Garti, N. Release of Markers from the Inner Water Phase of W / O / W Emulsions Stabilized by Silicone Based Polymeric Surfactants. J. Controlled Release 1995, 33 (1), 1–12. 10.1016/0168-3659(94)00029-T.

(70) Ladeveze, S.; Zurek, P. J.; Kaminski, T. S.; Emond, S.; Hollfelder, F. Versatile Product Detection via Coupled Assays for Ultrahigh-Throughput Screening of Carbohydrate-Active Enzymes in Microfluidic Droplets. ACS Catal. 2023, 13 (15), 10232–10243. 10.1021/acscatal.3c01609.

(71) Medcalf, E. J.; Gantz, M.; Kaminski, T. S.; Hollfelder, F. Ultra-High-Throughput Absorbance-Activated Droplet Sorting for Enzyme Screening at Kilohertz Frequencies. Anal. Chem. 2023, 95 (10), 4597–4604. 10.1021/acs.analchem.2c04144.

(72) Baret, J.-C.; Miller, O. J.; Taly, V.; Ryckelynck, M.; El-Harrak, A.; Frenz, L.; Rick, C.; Samuels, M. L.; Hutchison, J. B.; Agresti, J. J.; Link, D. R.; Weitz, D. A.; Griffiths, A. D. Fluorescence-Activated Droplet Sorting (FADS): Efficient Microfluidic Cell Sorting Based on Enzymatic Activity. Lab. Chip 2009, 9 (13), 1850–1858. 10.1039/B902504A.

(73) Yan, R.; Wang, W.; Vuong, T. V.; Xiu, Y.; Skarina, T.; Di Leo, R.; Gatenholm, P.; Toriz, G.; Tenkanen, M.; Stogios, P. J.; Master, E. R. Structural Characterization of the Family GH115 α-Glucuronidase from Amphibacillus Xylanus Yields Insight into Its Coordinated Action with α-Arabinofuranosidases. New Biotechnol. 2021, 62, 49–56. 10.1016/j.nbt.2021.01.005.

(74) Kolenová, K.; Ryabova, O.; Vršanská, M.; Biely, P. Inverting Character of Family GH115 α-Glucuronidases. FEBS Lett. 2010, 584 (18), 4063–4068. 10.1016/j.febslet.2010.08.031.

(75) Rogowski, A.; Baslé, A.; Farinas, C. S.; Solovyova, A.; Mortimer, J. C.; Dupree, P.; Gilbert, H. J.; Bolam, D. N. Evidence That GH115 α-Glucuronidase Activity, Which Is Required to Degrade Plant Biomass, Is Dependent on Conformational Flexibility *. J. Biol. Chem. 2014, 289 (1), 53–64. 10.1074/jbc.M113.525295.

(76) MacMillan, S.; Hosgood, S. A.; Walker-Panse, L.; Rahfeld, P.; Macdonald, S. S.; Kizhakkedathu, J. N.; Withers, S. G.; Nicholson, M. L. Enzymatic Conversion of Human Blood Group A Kidneys to Universal Blood Group O. Nat. Commun. 2024, 15 (1), 2795. 10.1038/s41467-024-47131-9.

(77) Ezeilo, U. R.; Zakaria, I. I.; Huyop, F.; Wahab, R. A. Enzymatic Breakdown of Lignocellulosic Biomass: The Role of Glycosyl Hydrolases and Lytic Polysaccharide Monooxygenases. Biotechnol. Biotechnol. Equip. 2017, 31 (4), 647–662. 10.1080/13102818.2017.1330124.

(78) Vuong, T. V.; Master, E. R. Enzymatic Production of 4-O-Methyl d-Glucaric Acid from Hardwood Xylan. Biotechnol. Biofuels 2020, 13 (1), 51. 10.1186/s13068-020-01691-2.

(79) Hu, H.; Li, J.; Jiang, W.; Jiang, Y.; Wan, Y.; Wang, Y.; Xin, F.; Zhang, W. Strategies for the Biological Synthesis of D-Glucuronic Acid and Its Derivatives. World J. Microbiol. Biotechnol. 2024, 40 (3), 94. 10.1007/s11274-024-03900-8.

(80) Yu, J.; Qian, Z.; Zhong, J. Advances in Bio-based Production of Dicarboxylic Acids Longer than C4. Eng. Life Sci. 2018, 18 (9), 668–681. 10.1002/elsc.201800023.

(81) Liguori, R.; Ventorino, V.; Pepe, O.; Faraco, V. Bioreactors for Lignocellulose Conversion into Fermentable Sugars for Production of High Added Value Products. Appl. Microbiol. Biotechnol. 2016, 100 (2), 597–611. 10.1007/s00253-015-7125-9.

(82) Fischlechner, M.; Schaerli, Y.; Mohamed, M. F.; Patil, S.; Abell, C.; Hollfelder, F. Evolution of Enzyme Catalysts Caged in Biomimetic Gel-Shell Beads. Nat. Chem. 2014, 6 (9), 791–796. 10.1038/nchem.1996.

(83) Kohler, T. N.; De Jonghe, J.; Ellermann, A. L.; Yanagida, A.; Herger, M.; Slatery, E. M.; Weberling, A.; Munger, C.; Fischer, K.; Mulas, C.; Winkel, A.; Ross, C.; Bergmann, S.; Franze, K.; Chalut, K.; Nichols, J.; Boroviak, T. E.; Hollfelder, F. Plakoglobin Is a Mechanoresponsive Regulator of Naive Pluripotency. Nat. Commun. 2023, 14 (1), 4022. 10.1038/s41467-023-39515-0.

(84) Fryer, T.; Rogers, J. D.; Mellor, C.; Kohler, T. N.; Minter, R.; Hollfelder, F. Gigavalent Display of Proteins on Monodisperse Polyacrylamide Hydrogels as a Versatile Modular Platform for Functional Assays and Protein Engineering. ACS Cent. Sci. 2022, 8 (8), 1182–1195. 10.1021/acscentsci.2c00576.

(85) Zinchenko, A.; Devenish, S. R. A.; Kintses, B.; Colin, P.-Y.; Fischlechner, M.; Hollfelder, F. One in a Million: Flow Cytometric Sorting of Single Cell-Lysate Assays in Monodisperse Picolitre Double Emulsion Droplets for Directed Evolution. Anal. Chem. 2014, 86 (5), 2526–2533. 10.1021/ac403585p.

(86) Tauzin, A. S.; Pereira, M. R.; Van Vliet, L. D.; Colin, P.-Y.; Laville, E.; Esque, J.; Laguerre, S.; Henrissat, B.; Terrapon, N.; Lombard, V.; Leclerc, M.; Doré, J.; Hollfelder, F.; Potocki-Veronese, G. Investigating Host-Microbiome Interactions by Droplet Based Microfluidics. Microbiome 2020, 8 (1), 141. 10.1186/s40168-020-00911-z.

(87) Simon-Baram, H.; Roth, S.; Niedermayer, C.; Huber, P.; Speck, M.; Diener, J.; Richter, M.; Bershtein, S. A High-Throughput Continuous Spectroscopic Assay to Measure the Activity of Natural Product Methyltransferases. ChemBioChem 2022, 23 (17), e202200162. 10.1002/cbic.202200162.

(88) Chouhan, B. P. S.; Maimaiti, S.; Gade, M.; Laurino, P. Rossmann-Fold Methyltransferases: Taking a “β-Turn” around Their Cofactor, S-Adenosylmethionine. Biochemistry 2019, 58 (3), 166–170. 10.1021/acs.biochem.8b00994.

(89) Radnai, L.; Stremel, R. F.; Sellers, J. R.; Rumbaugh, G.; Miller, C. A. A Semi High-Throughput Adaptation of the NADH-Coupled ATPase Assay for Screening of Small Molecule ATPase Inhibitors. J. Vis. Exp. JoVE 2019, No. 150, 10.3791/60017. https://doi.org/10.3791/60017.

(90) McFarlane, C. R.; Murray, J. W. A Sensitive Coupled Enzyme Assay for Measuring Kinase and ATPase Kinetics Using ADP-Specific Hexokinase. Bio-Protoc. 2020, 10 (9), e3599. 10.21769/BioProtoc.3599.

(91) Chiku, T.; Pullela, P. K.; Sem, D. S. A Dithio-Coupled Kinase and ATPase Assay. J. Biomol. Screen. 2006, 11 (7), 844–853. 10.1177/1087057106292142.

(92) Kaltwasser, H.; Schlegel, H. G. NADH-Dependent Coupled Enzyme Assay for Urease and Other Ammonia-Producing Systems. Anal. Biochem. 1966, 16 (1), 132–138. 10.1016/0003-2697(66)90088-1.

(93) Smith, H. Q.; Li, C.; Stanley, C. A.; Smith, T. J. Glutamate Dehydrogenase, a Complex Enzyme at a Crucial Metabolic Branch Point. Neurochem. Res. 2019, 44 (1), 117–132. 10.1007/s11064-017-2428-0.

(94) Relyea, H. A.; van der Donk, W. A. Mechanism and Applications of Phosphite Dehydrogenase. Bioorganic Chem. 2005, 33 (3), 171–189. 10.1016/j.bioorg.2005.01.003.

(95) Glieder, A.; Farinas, E. T.; Arnold, F. H. Laboratory Evolution of a Soluble, Self-Sufficient, Highly Active Alkane Hydroxylase. Nat. Biotechnol. 2002, 20 (11), 1135–1139. 10.1038/nbt744.

(96) de Miranda, A. S.; Milagre, C. D. F.; Hollmann, F. Alcohol Dehydrogenases as Catalysts in Organic Synthesis. Front. Catal. 2022, 2.

(97) Kroutil, W.; Mang, H.; Edegger, K.; Faber, K. Recent Advances in the Biocatalytic Reduction of Ketones and Oxidation of Sec-Alcohols. Curr. Opin. Chem. Biol. 2004, 8 (2), 120–126. 10.1016/j.cbpa.2004.02.005.

(98) Woodward, J.; Mattingly, S. M.; Danson, M.; Hough, D.; Ward, N.; Adams, M. In Vitro Hydrogen Production by Glucose Dehydrogenase and Hydrogenase. Nat. Biotechnol. 1996, 14 (7), 872–874. 10.1038/nbt0796-872.

(99) Khersonsky, O.; Röthlisberger, D.; Wollacott, A. M.; Murphy, P.; Dym, O.; Albeck, S.; Kiss, G.; Houk, K. N.; Baker, D.; Tawfik, D. S. Optimization of the In-Silico-Designed Kemp Eliminase KE70 by Computational Design and Directed Evolution. J. Mol. Biol. 2011, 407 (3), 391–412. 10.1016/j.jmb.2011.01.041.

(100) Khersonsky, O.; Röthlisberger, D.; Dym, O.; Albeck, S.; Jackson, C. J.; Baker, D.; Tawfik, D. S. Evolutionary Optimization of Computationally Designed Enzymes: Kemp Eliminases of the KE07 Series. J. Mol. Biol. 2010, 396 (4), 1025–1042. 10.1016/j.jmb.2009.12.031.

(101) Röthlisberger, D.; Khersonsky, O.; Wollacott, A. M.; Jiang, L.; DeChancie, J.; Betker, J.; Gallaher, J. L.; Althoff, E. A.; Zanghellini, A.; Dym, O.; Albeck, S.; Houk, K. N.; Tawfik, D. S.; Baker, D. Kemp Elimination Catalysts by Computational Enzyme Design. Nature 2008, 453 (7192), 190–195. 10.1038/nature06879.

(102) Jensen, R. A. Enzyme Recruitment in Evolution of New Function. Annu. Rev. Microbiol. 1976, 30, 409–425. 10.1146/annurev.mi.30.100176.002205.

(103) O’Brien, P. J.; Herschlag, D. Catalytic Promiscuity and the Evolution of New Enzymatic Activities. Chem. Biol. 1999, 6 (4), R91–R105. 10.1016/S1074-5521(99)80033-7.

(104) Schloss, P. D.; Handelsman, J. Biotechnological Prospects from Metagenomics. Curr. Opin. Biotechnol. 2003, 14 (3), 303–310. 10.1016/s0958-1669(03)00067-3.

